# A Genome-wide Survey of Mutations in the Jurkat Cell Line

**DOI:** 10.1101/118117

**Authors:** Louis Gioia, Azeem Siddique, Steven R. Head, Daniel R. Salomon, Andrew I. Su

**Author notes:** Deceased November 10, 2016 (Daniel R. Salomon).

## Abstract

The Jurkat cell line has an extensive history as a model of T cell signaling. But at the turn of the 21st century, some irregularities were observed in Jurkat’s expression of central regulators of T cell receptor signaling, which raised doubts about how closely the cell line paralleled normal human T cells. While numerous expression deficiencies have been described in Jurkat, genetic explanations have only been provided for a handful of defects. Here, we report a comprehensive catolog of genomic variation in the Jurkat cell line based on whole-genome sequencing. With this list of all detectable, non-reference sequences, we prioritize potentially damaging mutations by mining public databases for functional effects. We confirm the majority of documented mutations in Jurkat and propose links from detrimental gene variants to observed expression abnormalities in the cell line. This work ties together decades of molecular experiments and serves as a resource that will streamline both the interpretation of past research and the design of future Jurkat studies.

## Introduction

The Jurkat cell line was isolated in 1977 from the blood of a fourteen-year-old boy with Acute Lymphoblastic Leukemia (1). It was one of the first *in vitro* systems for studying T-cell biology and helped to produce an incredible number of discoveries and publications (Fig. 1) (2).

**Figure 1:**
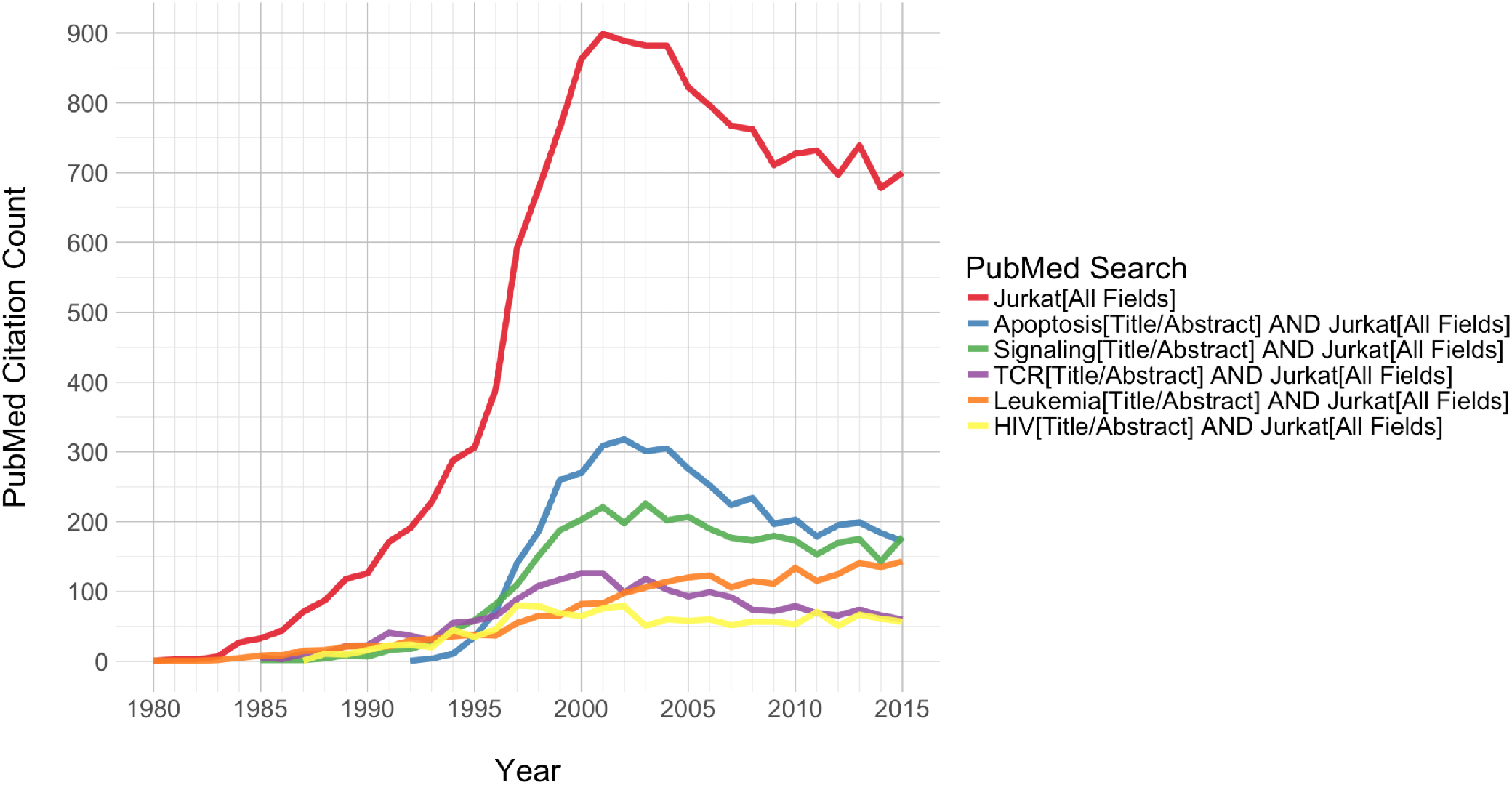
Jurkat publication trends. Yearly publication counts for PubMed queries. Representative queries are given in the legend. Note that these query descriptions are abbreviations of more detailed search terms, which are provided in the Methods section.

As the workhorse behind a diverse array of molecular investigations, the Jurkat cell line revealed the foundations for our modern understanding of multiple signaling pathways. Most notably, studies of Jurkat cells established the bulk of what is currently known about T-cell receptor (TCR) signaling (2). However, at the turn of the 21st century, as the use of Jurkat as a model T-cell line was reaching its height, some abnormalities in the cell line began to come to light.

Problems were first noticed in the form of gene expression defects. The most publicized of these defects was aberrant PI3K signaling due to the absence of PTEN and INPP5D (SHIP) in Jurkat cells (2). The loss of these two central regulators of phosphatidylinositol signaling was proposed as the cause of the previously-documented, constitutive activation of PI3K signaling, a major mediator of downstream TCR signaling events (3). This fundamental TCR signaling defect in Jurkat led many researchers to question its validity as a model system for T-cell studies (2). Although the number of publications using Jurkat dropped off over the following decade, it is still widely used in biomedical research (Figure 1).

Defect detection up to now has been primarily based on top-down approaches, requiring knowledge of signaling or expression defects, which leads to interrogations of specific coding sequences. While multiple genetic defects have been described over the past few decades, these top-down approaches are limited in scope and have failed to provide a broader understanding of Jurkat biology.

Modern sequencing technology allows for interrogation of the entire genome. In contrast to top-down techniques, whole-genome sequencing (WGS) allows us to investigate genetic defects from the bottom up, with the potential to extend our understanding of abnormalities in Jurkat. Thus, in this study, we used shotgun sequencing to perform genome-scale characterization of genomic variants in this commonly-used cell line.

## Results

### Sequencing and Variant Callers

Whole-genome sequencing of the Jurkat cell line produced over 366 million 100bp paired-end reads and over 531 million 150bp paired-end reads, totaling over 116 billion sequenced bases. More than 98% of the reads were successfully aligned to the hg19 human reference genome with the Burrows-Wheeler Aligner (4), totaling over 110 billion aligned bases. This gave an average coverage of ~36x across the hg19 reference sequence, with over 10x depth of coverage for 78.8% of the genome. The aligned reads were then used to detect both small and large genomic variants in the Jurkat genome.

In order to utilize all of the information available in the WGS data, we employed a suite of variant calling tools for the identification of all major types of genomic variants. Each tool uses a certain type of sequence information to identify specific categories of variants. Our variant caller suite consisted of four distinct tools and algorithms: The Genome Analysis Toolkit, Pindel, BreakDancer, and CNVnator *(5)–(8).*

The Genome Analysis Toolkit (GATK) from the Broad Institute uses De Bruijn graph-based models to identify single-nucleotide substitutions and small insertions and deletions. Pindel’s split-read approach can also detect small insertions and deletions, as well as inversions, tandem duplications, and inter-chromosomal translocations. BreakDancer compares the distance between aligned read pairs to the insert size distribution from the sequencing library in order to find large structural variants. CNVnator uses read-depth information and a mean-shift algorithm to assign copy number levels across the genome and identify deletion and duplication events.

In order for GATK to call small variants, it must be told how many alleles to expect at each position. As such, an accurate estimate of Jurkat ploidy is required before GATK can be used. While both the original 1977 publication and the American Type Culture Collection (ATCC) report that Jurkat is diploid (9), other publications refute this description. The first karyotypes of the Jurkat cell line were published by Snow and Judd in 1987, who found that Jurkat was hypotetraploid (10). A few years later, tetraploidy was corroborated by an investigation of p53 mutations, which found that the Jurkat cell line contained 4 separate p53 alleles (11). More recent reports confirm Jurkat tetraploidy. The German Collection of Microorganisms and Cell Cultures (DSMZ) describes the Jurkat karyotype as a “human flat-moded hypotetraploid karyotype with 7.8% polyploidy” (12). In addition, a multicolor-Fluorescence In Situ Hybridization study from 2013 found within-culture mosaicism on a tetraploid background (13).

### Variant Calls

Given the previous reports of tetraploidy, we ran GATK with a ploidy count of 4. GATK identified nearly 5 million variants, comprising ~3.5 million single-nucleotide substitutions, ~1.0 million small deletions, and ~357 thousand small insertions, across over 4.6 million variant loci. Basic metrics for the GATK variant calls are consistent with normal human samples. The ratio of homozygous to heterozygous variant loci is 0.635, which is in the range of previously reported ratios, and the ratio of transitions to transversions is 2.10, which is the expected value for human genomes (14). The number of single-nucleotide substitutions is similar to previously reported values. However, the total number of indels is higher than published values from human WGS studies, which generally detect fewer than 700 thousand indels by shotgun sequencing (14). To date, the highest number of indels identified in a single human genome was ~850 thousand—determined via Sanger sequencing of J.C. Venter’s genome (15). This enrichment for indels, especially deletions, in Jurkat is likely to be at least partially due to the redundancy of the tetraploid genome.

The Pindel variant caller detected 1.4 million deletions, 740 thousand insertions, 18 thousand duplications, 150 thousand inversions, and 4 inter-chromosomal translocations. The split-read approach is markedly similar to GATK’s method for the detection of small insertions and deletions. GATK also uses split-reads, but its detection of variants relies on an assembly-based method that is limited to small sequence differences between the reads and the reference genome. Accordingly, the small indels called by both methods should be similar. As expected, in the Jurkat call set, over 85% of the deletions and over 65% of the insertions that were identified by GATK have direct matches in the Pindel calls.

BreakDancer identified 6,128 deletions, 18 insertions, 183 inversions, 1,981 intra-chromosomal translocations, and 113 inter-chromosomal translocations.

CNV calls from CNVnator are presented in Figure 2 by percentage of the genome. A plot of the raw read depth density is provided in Supplementary Figure S1. CNVnator reports a modal copy number of 4 in Jurkat, representing over 65% of the genome and corroborating reports of tetraploidy. From the CNVnator results, we identified 2,499 deletion sites (CN ≤ 1), of which 218 were homozygous (CN = 0), and 1,863 duplication sites (CN ≤ 5).

**Figure 2:**
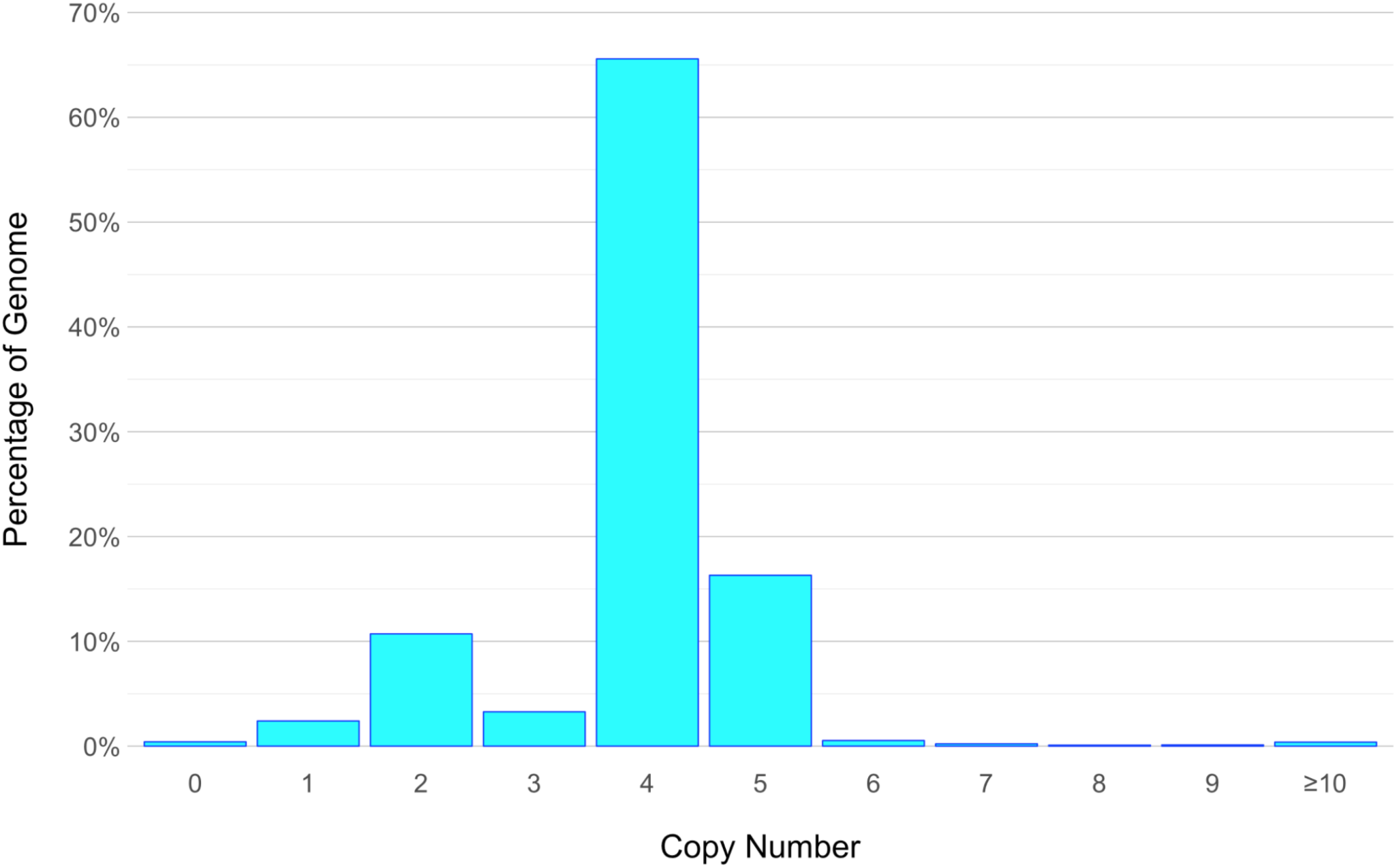
Histogram of DNA copy number in Jurkat. Binned copy number alterations as fractions of the genome.

The structural variant calls from each tool were compared and merged with specific considerations made for each category of variant and each detection tool (see Methods). Short and long insertions and deletions were defined using a cutoff of 50bp, in accordance with the structural variant databases from NCBI (16). The numbers of variants called by each tool, along with the proportion of overlapping loci and total number of merged calls, are provided in Table 1.

**Table 1.** Variant loci counts from each tool. The percentage of sites that overlap the other tools is provided where applicable

|  | **GATK** | **Pindel** | **BreakDancer** | **CNVnator** | **Merged** |
| --- | --- | --- | --- | --- | --- |
| **Substitutions** | 3,520,988 |  |  |  | 3,520,988 |
| **Short Hom. Deletion** | 170,397 |  |  |  | 170,397 |
| **Short Deletion** | 841,001<br>(70%) | 1,239,299<br>(47%) | 47<br>(0%) |  | 1,460,321 |
| **Short Insertion** | 326,446<br>(65%) | 616,298<br>(35%) |  |  | 729,727 |
| **Long Hom. Deletion** | 326<br>(0%) |  |  | 108<br>(0%) | 434 |
| **Long Deletion** | 1,904<br>(61%) | 118,610<br>(1.4%) | 6,081<br>(10%) | 2,499<br>(1.2%) | 125,397 |
| **Long Insertion** | 1,039<br>(31%) | 125,918<br>(0.25%) | 18<br>(0%) |  | 126,657 |
| **Duplication** |  | 17,762<br>(22%) |  | 1,863<br>(24%) | 15,288 |
| **Inversion** |  | 149,545<br>(0.0087%) | 183<br>(7.1%) |  | 149,715 |
| **Intra. Translocation** |  |  | 1,981 |  | 1,981 |
| **Inter. Translocation** |  | 4<br>(0%) | 113<br>(0%) |  | 117 |

Most types of variants were called by multiple tools. However, the number of variants called by each tool and the number of variant calls that were unique to each tool varied greatly between variant classes and individual variant callers (Table 1). Furthermore, each tool differed in the sizes of variants that it called (Figures S2-S8).

The relative contributions of each variant caller to the total set of merged calls are displayed in Figure 3. Pindel calls dominated the merged variant sets, with the exception of translocations. This unmatched number of Pindel calls can be attributed to the power of the split-read approach. On the other hand, Pindel calls are limited in their utility due to the tool’s inability to determine allele frequencies.

**Figure 3:**
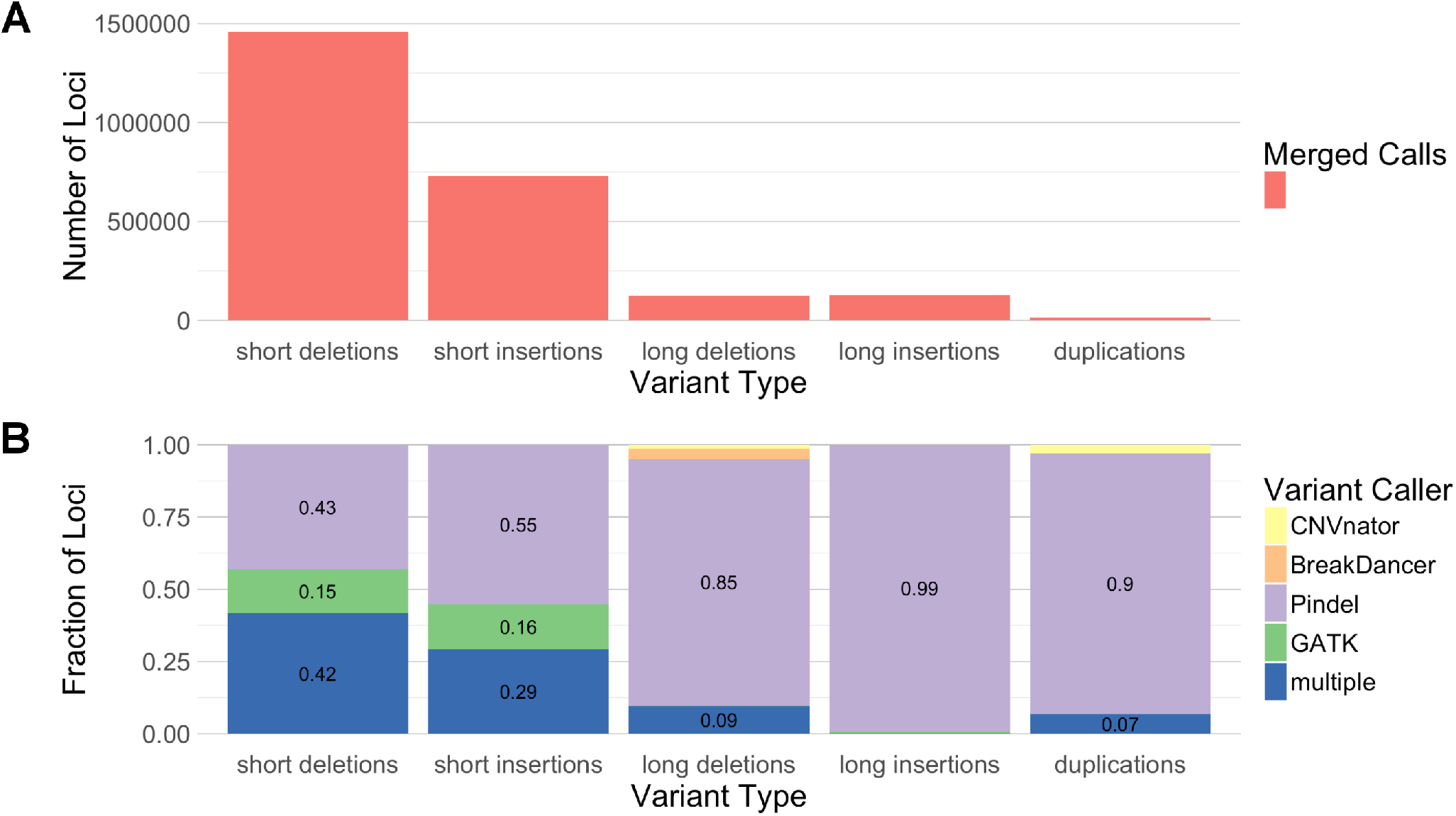
Comparison of numbers of variant loci called by each tool. (**A**) Total number of merged variant loci called by all tools for different variant types. (**B**) Fraction of merged variant loci called by each tool for different variant types.

In contrast to Pindel’s detection power and lack of allele annotations, GATK and CNVnator are both limited in the range of variant sizes that they can detect but are able to consider all alleles. Therefore, while Pindel calls make up the majority of detected variants, GATK and CNVnator calls were prioritized in our investigations of variant consequence.

### Comparisons to Databases

After creating the merged variant sets, we compared them to databases of previously identified variants in order to assess the novelty of the genomic variants that were detected in Jurkat. We used dbSNP and DGV as resources for known short and long variants, respectively (17), (18). Both of these databases contain the variants that were identified by the 1000Genomes project in addition to variants cataloged by other sources. Comparisons of short variants—including single-nucleotide substitutions, short deletions, and short insertions—to variants found in the 1000Genomes project and dbSNP are given in Figure 4. Single-nucleotide substitutions showed the greatest number of matches, while fewer than half of the short insertions and deletions were found in dbSNP. An even greater reduction in the number of database matches was seen in the long structural variant database comparisons (Figure 4). The differences in the number of database matches between single-nucleotide variants, short indels, and long structural variants are likely due to several factors. The feasibility of structural variant detection, combined with the paucity of studies investigating these larger variants, are major contributors to these differences, but the increased mutational sample space of larger variants may also play a role.

**Figure 4:**
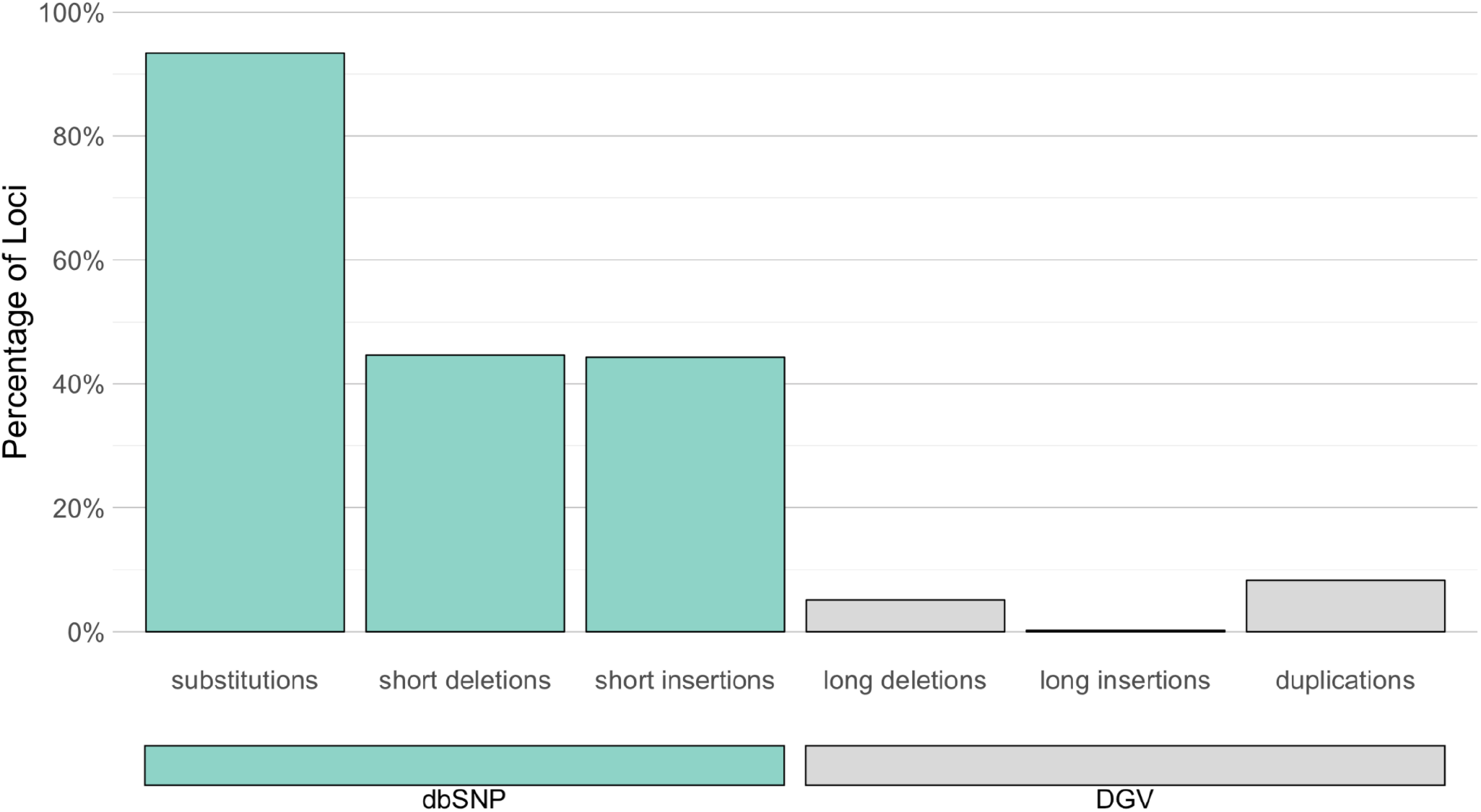
Jurkat variants with database matches. Jurkat variants loci that have matches in dbSNP (short variants) and DGV (long variants) as percentage of total Jurkat variant sites for each type of variant. Number of databases matches over the number of Jurkat variant loci: 3.29M / 3.52M substitutions; 652K / 1.46M short deletions; 323K / 730K short insertions; 6.38K / 125K long deletions; 286 / 127K long insertions; 1.27K / 15.3K duplications.

We also compared our SNV and small indel calls to those found in Jurkat by the COSMIC Cell Line project (19). Our WGS approach identified nearly 10x as many SNVs as were detected by COSMIC via microarray. However, of the ~408 thousand Jurkat SNVs in COSMIC, we uncovered over 383 thousand (94%) matching single-nucleotide variants. Within the matching SNV calls, genotypes between the two call sets agreed at over 97% of loci. The same level of agreement was observed for both the ~174 thousand homozygous COSMIC calls and the ~210 thousand heterozygous COSMIC calls. Deletion and insertion calls showed less overlap, but we were able to find 67% of the 18 thousand COSMIC deletion calls and 40% of the 2,260 COSMIC insertion calls in our data.

Our final comparison to previously identified variants focused on rare, pathogenic variants from the ClinVar database. After removing records without assertion criteria, corresponding to a review status of zero stars, 10 Jurkat variants were reported as pathogenic by ClinVar (Table 2). Interestingly, 6 of the 10 variants, involving 5 separate genes, are thought to cause cancer. The other pathogenic ClinVar matches are associated with severe developmental defects. Long deletions and duplications from Jurkat were also found in ClinVar, but the annotations do not contain gene information and are generally less informative (Supplementary Tables 1 & 2).

**Table 2:** Jurkat variants found in the ClinVar database.

| rsID | Jurkat AF | Gene | Phenotype | ClinVar Accession |
| --- | --- | --- | --- | --- |
| ***ClinVar Substitutions*** |  |  |  |  |
| rs63750636 | 1.0 | MSH2 | Lynch syndrome | RCV000076405.3 |
| rs397517342 | 0.75 | CDH23 | Usher syndrome type 1D | RCV000039224.2 |
| rs397516435 | 0.25 | TP53 | Li-Fraumeni syndrome | RCV000205265.3 |
| ***ClinVar Short Deletions*** |  |  |  |  |
| rs63750075 | — | MSH6 | Lynch syndrome | RCV000074711.2 |
| rs398122841 | — | BAX | Carcinoma of colon | RCV000010120.5 |
| rs397508104 | — | KCNQ1-(AS1) | Long QT syndrome | RCV000046039.3 |
| rs750664956 | — | ASPM | Not provided | RCV000217980.1 |
| rs786204835 | — | PURA | Not provided | RCV000169739.5 |
| ***ClinVar Short Insertions*** |  |  |  |  |
| rs397507178 | — | RAD50 | Hereditary cancer | RCV000030958.3 |
| rs398122840 | — | BAX | Carcinoma of colon | RCV000010119.5 |

Moving from established to predicted effects, we used SnpEff to predict the functional consequences of the GATK-called small variants. SnpEff identified 9,997 synonymous and 10,984 nonsynonymous mutations. Among the nonsynonymous mutations, 252 variants are nonsense mutations and 10,732 variants are missense mutations. ‘High Impact’ functional effects were predicted for 1141 of the small variant loci, of which 747 variants were determined to be rare (MAF < 0.001) in the Exome Aggregation Consortium (ExAC) dataset of over 60 thousand human samples (20). These rare, high-impact variants were predicted to affect 678 genes.

A second set of ‘High Impact’ variants was created from the homozygous deletion calls that intersected coding exons. This high-impact, homozygous deletion set includes 120 variant loci across 129 genes.

All sets of variants, including those of high impact, appear to be distributed across the genome (Figure 5). However, even if the mutations are randomly distributed, it is still possible that some biological processes are more affected than others. The two sets of highly impacted genes were combined, producing a set of 781 unique genes. This list of likely damaged genes was used to probe selected gene set databases from MSigDB (21). The top 5 enriched gene sets are displayed in table 3.

**Figure 5:**
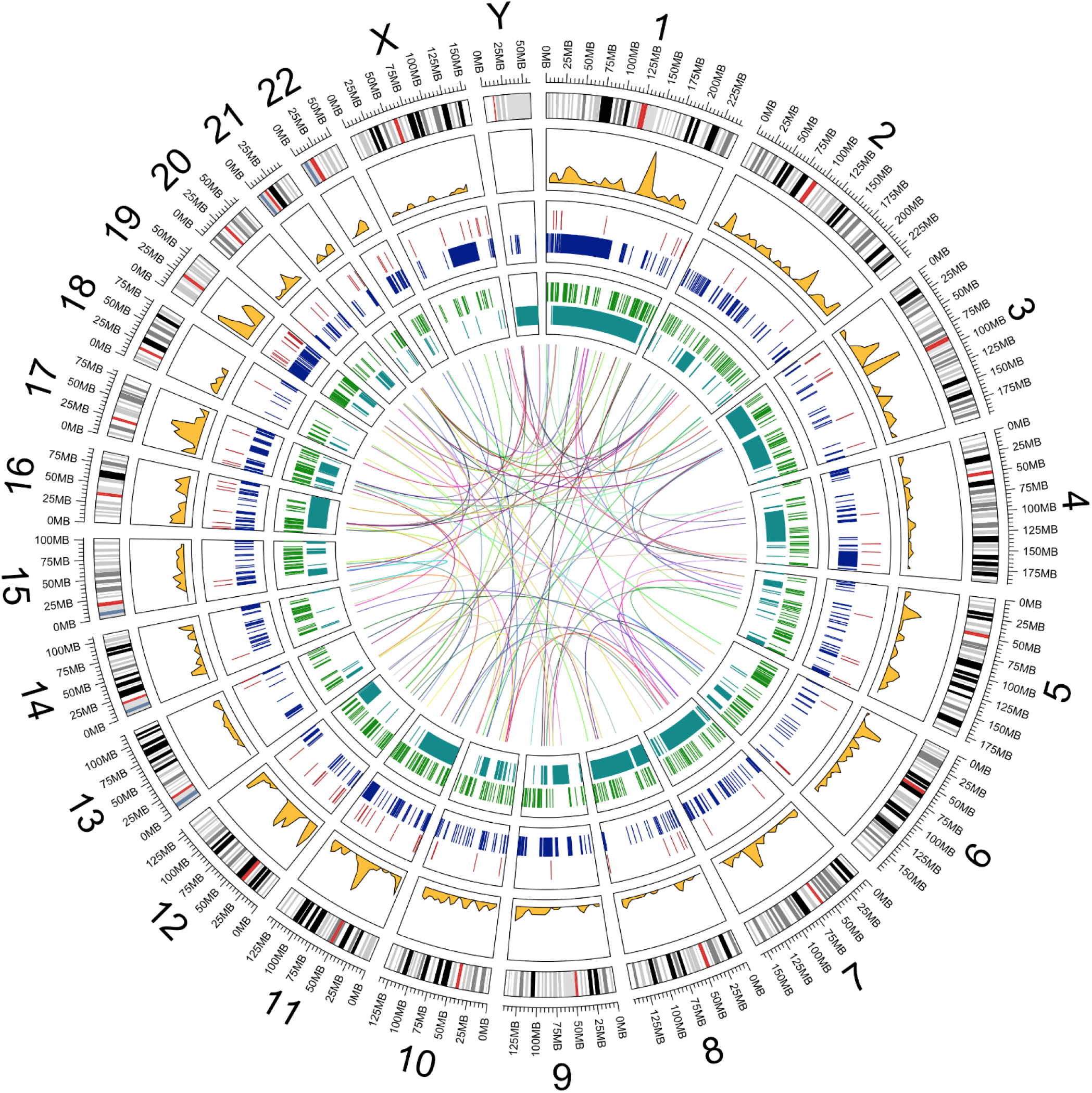
Genomic variation distributions. Distributions of multiple types of variants across the Jurkat genome. Plotted data listed from outside-in: 1. hg19 genome ideogram (gray); 2. Density of SnpEff “High Impact” SNVs with rare ExAC allele frequencies (gold); 3. Homozygous deletions that lie in coding exons (red); 4. Deletions longer than 25kb (blue); 5. Insertions longer than 50bp that lie in coding exons (green); 6. Inversions longer than 25kb (cyan); 7. Interchromosomal translocations (center).

**Table 3:** Gene sets enriched for highly impacted genes.

|  |
| --- |
| **GO: CHROMOSOME ORGANIZATION**<br>*Overlap: 51/1009 p-value= 0.00011*<br>**Genes:** APBB1, ATXN3, ATXN7, BAZ2A, BCORL1, BRD8, CDC14A, CDCA5, CDYL, CENPT, CLASP1, CREBBP, EHMT1, GATAD2B, GTF2H3, HDAC4, KAT2A, KDM5B, KIF23, KNTC1, MLH3, MSH2, NCAPD2, NDC80, NOC2L, PIBF1, PRIM2, PTGES3, RAD50, RBL2, RSF1, SETX, SLX4, SMARCC2, SMC3, TCF7L2, TEP1, TET1, TEX14, TOP1MT, TP53, TTN, USP15, VPRBP, YEATS2, ZNF304, ZNF462 |
| **GO: CELL CYCLE**<br>*Overlap: 62/1316 p-value= 0.00015*<br>**Genes:** ADCY3, ANAPC5, APBB1, ARHGEF2, BAX, CDC14A, CDC27, CDCA5, CDK14, CECR2, CENPT, CEP164, CKAP2, CLASP1, DYNC1H1, DYNC1I2, FANCI, HINFP, HSP90AA1, INTS3, IQGAP3, KIAA0430, KIAA1377, KIF23, KNTC1, KRT18, MACF1, MAP3K8, MAP9, MCM8, MKI67, MLH3, MNS1, MSH2, NCAPD2, NDC80, NUP214, NUP98, OFD1, ORC1, PHLDA1, PIBF1, PRIM2, PSMD3, PTEN, PYHIN1, RAD50, RBL2, RUVBL1, SMC3, SON, TCF7L2, TEX14, THAP1, TP53, TP53BP1, TPR, TRIOBP, TSC1, TTK, TTN, ZFHX3 |
| **GO: CARBOHYDRATE DERIVATIVE BIOSYNTHETIC PROCESS**<br>*Overlap: 34/595 p-value= 0.00017*<br>**Genes:** ADCY2, ADCY3, ADCY9, ALG1L2, ALG9, B3GALT1, B3GNT6, BCAN, BMPR2, C1GALT1C1, CANT1, CHST15, CHSY1, GAL3ST4, GPC6, GUCY2C, GXYLT1, HAS3, KIAA2018, MUC16, MUC19, MUC3A, MUC6, NDST4, OMD, PHLDA1, PIGS, PRKCSH, SLC25A13, ST3GAL3, ST3GAL5, TET1, UGCG, UGP2 |
| **GO: NEGATIVE REGULATION OF ORGANELLE ORGANIZATION**<br>*Overlap: 25/387 p-value= 0.00020*<br>**Genes:** ARHGEF2, CDC14A, CKAP2, CLASP1, DYNC1H1, KIAA1377, KIF23, LIMA1, MAP9, MSH2, NDC80, NOC2L, OFD1, OTUB1, PIBF1, RAD50, SMC3, SPTA1, SPTAN1, TET1, TEX14, TPR, TRIOBP, TTK, UBQLN4 |
| **GO: IMMUNE SYSTEM PROCESS**<br>*Overlap: 85/1984 p-value= 0.00023*<br>**Genes:** ABCB5, ADAM17, AGBL5, AIM2, AP3B1, APOB, ARHGEF2, BAX, C1orf177, C7, CCL13, CD14, CD177, CEACAM8, CLNK, CREBBP, CTLA4, CYFIP2, DEFB126, DHX58, DYNC1H1, DYNC1I2, DYNC2H1, ENDOU, ENPP3, F2, FN1, HDAC4, HLA-DRB5, HNRNPK, HSH2D, HSP90AA1, IGJ, IL10RB, IL27RA, IL2RG, ILF2, INPP5D, IPO7, ITGA6, KIF23, KIF3C, KIR2DS4, KLC2, LILRA3, MAP3K1, MAP3K8, MSH2, NCAM1, NLRC3, NLRC5, OAS1, OTUB1, PAPD4, PIBF1, PODXL, PRKACG, PSMD3, PTEN, RHOH, SAMHD1, SARM1, SEC31A, SECTM1, SHC1, SLC3A2, SLFN11, SPEF2, SPTA1, STAT5B, SYK, SYNCRIP, TAB2, TAPBP, TEK, TMIGD2, TNFSF4, TNK2, TRIL, TRIM10, TSC1, ULBP1, VPRBP, WDR7, WIPF1 |

As might be expected from a cancer cell line, the damaged genes in Jurkat are involved in genome, cell cycle, and cytoskeleton maintenance, as well as sugar processing. The enrichment of damaged genes that are involved in the immune system is particularly interesting given the Jurkat cell line’s role in establishing our current understanding of T-cell immune responses.

While the gene set enrichment analysis aided in categorizing the many genetic aberrations in the Jurkat cell line, most of the top-enriched sets are broad, suggesting gross defects across general biological processes. These findings reinforce the growing body of literature that has cataloged numerous irregularities in Jurkat biology, but they also imply that the deviation from normal T-cell biology may be more extensive than previous studies had reported.

### Defective Pathways

By leveraging the deep history of the Jurkat cell line, in combination with our pathogenic and high-impact variant lists, we have distinguished three core pathways that are defective due to genomic aberrations in Jurkat— namely TCR signaling, genome stability, and O-linked glycosylation. This analysis is not exhaustive. Rather, we focused on pathways are well-supported by both the literature and our genomic analysis.

### TCR Signaling

The damaged genes affecting T-cell receptor signaling are PTEN, INPP5D, CTLA4, and SYK. TCR signaling in Jurkat was first called into question due to the lack of PTEN and INPP5D expression (3), (22). Both PTEN and INPP5D are lipid phosphatases that regulate PI3K signaling by degrading PtdIns(3,4,5)P3. PTEN mutations in Jurkat were first described by Sakai et al. in 1998. They found two separate alterations in exon 7 “without normal conformers present,” both of which introduced stop codons (23). We detected the same two heterozygous variants. SnpEff annotated one of these mutations as a frameshift variant and the other as a stop-gained variant, predicting that both of these variants would result in loss of function.

INPP5D (SHIP1) has long been known to not be expressed in the Jurkat cell line (3). We have identified a single-nucleotide substitution that changes codon 317 from glutamine to a stop codon, as well as a 47bp heterozygous deletion from hg19.chr2:234068130-234068177. These same mutations were detected in 2009 via targeted sequencing (24). Admittedly, the lack of allele resolution in our data precludes us from making definitive claims about these mutations, as we cannot distinguish which alleles were affected. Fortunately, the targeted sequencing study found the stop codon on one allele and the 47bp deletion on the others, both of which should block the production of a full length INPP5D transcript.

CTLA4 is a CD28 homolog that transmits an inhibitory signal to T cells. In 1993, Lindsten et al. noticed that “CTLA4 mRNA is not expressed nor induced in the Jurkat T cell line” (25). However, the reason for this lack of CTLA4 induction has not been proposed. More recent investigations have detected both the protein and the transcript, although the transcript was less abundant in Jurkat than in peripheral blood mononuclear cells (26). This finding seems to support the hypothesis that the CTLA4 protein is accumulated in the cytosol (26). Our analyses revealed a heterozygous, stop-gained, single-nucleotide substitution that converts codon 20 to a stop codon. This mutation was found in around half of the mapped reads and might be responsible for the decreased CTLA4 expression that has been observed in Jurkat cells, although other mechanisms may be at play.

SYK is a member of the Syk family of nonreceptor tyrosine kinases. It functions similarly to ZAP70 in transmitting signals from the T-cell receptor. In 1995, Fargnoli et al. reported that SYK is not expressed in the Jurkat cell line and contains a guanine insertion that causes a frameshift at codon 34. We identified the same heterozygous insertion in our sample, which is predicted to result in loss of function of the transcript, yet the mechanism behind the lack of expression of the other allele remains an open question.

Interestingly, Fargnoli et al. proposed that the lack of SYK expression in Jurkat “may have facilitated the initial identification and characterization of ZAP70 as the major Z-associated protein” (27). On the other hand, while the lack of SYK expression in Jurkat was subsequently confirmed, reconstitution studies suggest that SYK and ZAP70 occupy distinct roles in TCR signaling, with SYK displaying 100-fold greater kinase activity than ZAP70 *(28),(29).*

### Genome Stability

TP53, BAX, and MSH2 are all tumor suppressors involved in maintaining genomic stability that are severely mutated in Jurkat. The product of the TP53 gene is p53, which is a known deficiency in the Jurkat cell line (22). In 1990, Cheng and Haas detected a heterozygous, stop-gained single-nucleotide substitution in codon 196 (R196*) in Jurkat cells. They proposed that this mutation “may play a role in the genesis or in the tumorigenic progression of leukemic T cells” (11). We detected the same heterozygous mutation (rs397516435) in exon 6 of the TP53 gene and found that this mutation is associated with Li-Fraumeni syndrome (30), which is an autosomal dominant hereditary disorder that causes the early onset of tumors. This mutation is likely responsible for the consistent reports of p53 deficiencies in Jurkat cells.

While loss of p53’s protective effects is normally thought of as the mechanism behind tumorigenesis, in some cases, truncated p53 can gain oncogenic functions (31). Recent studies have revealed that stop-gained mutations in exon 6 of TP53 produce a truncated p53 isoform that seems to partially escape nonsense-mediated decay. These isoforms, termed p53*ψ*, lack canonical p53 transcriptional activity. Instead, they localize to the mitochondria, where they activate a pro-tumorigenic cellular program by regulating mitochondrial transition pore permeability through interaction with cyclophilin D (32). The Jurkat cell line’s expression of a p53 *ψ* isoform may contribute to the previously-reported, exaggerated Ca^2+^ release upon TCR activation (33).

BAX is a member of the Bcl-2 gene family and helps induce apoptosis. In the Jurkat cell line, BAX is not expressed due to the presence of two heterozygous frameshift mutations in codon 41 (34). All alleles are affected. We identified the same two variants, rs398122841 and rs398122840, each of which were found in approximately half of the mapped reads.

Investigations into microsatellite instability revealed that MSH2 is not expressed in Jurkat due to a stop-gained point mutation in exon 13 (35). We identified the same variant as a homozygous single-nucleotide substitution (rs63750636.)

MSH2 is involved in DNA mismatch repair, and this stop-gained variant is associated with hereditary nonpolyposis colorectal cancer (36).

### O-linked Glycosylation

The Jurkat cell line’s inability to properly synthesize O-glycans, due to deficient core 1 synthase, glycoprotein-N-acetylgalactosamine 3-beta-galactosyltransferase 1 (C1GALT1) activity, was first noticed in 1990 (37). This deficiency causes Jurkat to express the Tn antigen, which is associated with cancer and other pathologies. In 2002, Ju and Cummings reported that a single-nucleotide deletion in COSMC (C1GALT1C1), a chaperone for C1GALT1, was responsible for Jurkat’s truncated O-glycans. The deletion causes a frameshift and introduces a stop codon in the only exon of the COSMC gene. Ju and Cummings assumed that the Jurkat cell line had retained its diploid, male genome and possessed only one copy of the X chromosome. We now know that the Jurkat cell line has two copies of the X chromosome, but consistent with the original report, we have determined through deep sequencing that the mutation is, indeed, homozygous across Jurkat’s two X chromosomes (38).

## Discussion

We performed a bottom-up search for abnormalities in the Jurkat genome using short-read sequencing. We detect numerous examples of each examined variant type and use various strategies to tie these variants to functional effects. Our analysis identifies multiple dysfunctional pathways in the Jurkat cell line.

While some of the variants were previously detected using top-down methods, we were able to add hundreds of potentially damaging variants to the list of Jurkat’s genetic defects. Gene set enrichment analysis revealed that many of the affected genes lie in pathways that are commonly defective in cancer. The great number of potentially damaging genes, combined with the large-scale genomic rearrangements in Jurkat, make it difficult to pinpoint the cause of Jurkat’s biological abnormalities. However, some of the better-studied mutations, such as those reported as pathogenic by ClinVar, are likely to have significant effects on important signaling pathways.

In addition to these putatively damaging variants, we identified millions of mutations across all categories of genomic variants. The effects of these variants are less certain, but our comprehensive variant catalog will facilitate further investigations of the Jurkat genome and allow for re-analysis of Jurkat variants as more information about their effects becomes available.

Using our list of variants, we were also able to extensively search the literature for previously identified defects in Jurkat. We found many reports of the same variants that we had independently identified. Uncovering these publications required precise knowledge of damaged genes in Jurkat. They were difficult, if not impossible, to find using general queries and were published over a decade ago in a range of journals. Furthermore, with the exception of the PTEN and INPP5D defects, these reports had never been consolidated into a single resource, making our documentation of previous reports the first review of damaged genes in the Jurkat cell line.

The defects in these genes have the potential to confound prior findings in Jurkat, but the loss is unlikely to put a dent in the vast amount of knowledge that we have gained from this cell line. In fact, Jurkat’s expression deficiencies open the door for reconstitution experiments that, in other systems, would first require suppression of the gene products. Many studies have already put this idea into action. Transgenic expression of INPP5D and SYK constructs has already generated breakthroughs in our understanding of their biological activities (28,29,39,40).

Likewise, the presence of pathogenic protein isoforms may also facilitate new discoveries by enabling the study of these factors *in vitro,* without the introduction of transgenic systems. For example, our finding that Jurkat expresses p53*ψ*, a newly recognized p53 isoform with direct links to malignant transformation, suggests that the cell line could be used as a model system for examination of p53*ψ*’s non-canonical activities.

Our bottom-up analysis was mostly limited by the availability of variant effect information. We focused our analysis and discussion on the effects of short variants as much more is known about their functional consequences. While we were unable to derive much information from the larger structural variants that we detected, we hope that they will become more meaningful as the ability to detect and study structural variants increases in the future.

The possibility that further discoveries are present in our data still remains. To this end, we provide our full set of sequencing and variant data for future use. Sequencing data is available at the NCBI Sequence Read Archive (SRA) study SRP101994 (https://www.ncbi.nlm.nih.gov/sra/SRP101994). Raw variant caller output files, filtered and merged variant calls, and variant effect information are available at Zenodo (https://zenodo.org/record/400615) (41). The scripts used to produce this data are provided in a public repository at https://bitbucket.org/sulab/jurkat_variant_calling. Open access to our data will allow for reanalysis and will ease the discovery of further biological defects in the Jurkat cell line.

## Methods

### Sequencing

DNA library preparation was performed with the iGenomX RLP according to manufacturer’s specifications (http://www.igenomx.com/technology.html). Two lanes each of 100bp and 150bp paired-end sequencing were run on an Illumina HiSeq instrument, generating over 116 billion bases of whole-genome sequencing data.

### Data Preprocessing

Data quality was checked using the FastQC software version 0. 11.2 from Babraham Bioinformatics (http://www.bioinformatics.babraham.ac.uk/projects/fastqc/), which confirmed that the sequencing was of high quality. Each lane of sequencing data was aligned to the hg19 human reference genome using the bwa-mem algorithm from BWA version 0.7.10 with default options (4). Genome coverage was calculated with the genomecov command from the BEDtools software package version 2.25.0 (http://bedtools.readthedocs.io/en/stable/index.html), and average coverage was calculated as the number of mapped bases divided by the total number of bases in the reference genome. The Picard software toolkit version 1.103 from the Broad Institute (https://broadinstitute.github.io/picard/) was used to add read groups and mark duplicate reads in the bam files. Realignment around indels and recalibration of base quality scores were done with the Genome

Analysis Toolkit software version 3.3 from the Broad Institute (https://software.broadinstitute.org/gatk/). The 1000Genomes project phase 1 indels and the Mills and 1000Genomes gold standard indels were used as known sites for GATK’s IndelRealigner. Base quality score recalibration was performed using the same known indel sites plus the dbSNP build 137 known sites. The alignment files for each lane were then merged with Samtools version 0.1.19. Duplicate reads in the merged.bam file were marked before GATK variant calling, but this last step was skipped for the other variant callers.

### Variant Calling

Small variants were detected with GATK’s haplotype caller according to the recommended settings, i.e. the-stand_call_conf parameter was set to 30 and the-stand_emit_conf parameter was set to 10. The ploidy parameter was set to 4, corresponding to a tetraploid genome, which was assumed based on literature review. The GATK variant quality score recalibration (VQSR) scheme was run for both SNPs and indels, and only the last tranche was filtered out of the call set with the-ts_filter parameter set to 99.9. All of the recommended truth and training sets for VQSR were used with the recommended parameters for each resource.

After quality filtering, structural variant calls were extracted from the individual variant caller output files and converted to a BED file format for comparisons. Structural variant calls other than inter-chromosomal translocations were removed if they overlapped the centromeric or telomeric repeat regions of the hg19 reference. Overlaps were calculated with the BEDtools intersect command. Repeat regions were extracted from the UCSC Sequence and Annotation Downloads, hg19 gap database (http://hgdownload.cse.ucsc.edu/goldenPath/hg19/database/gap.txt.gz).

Pindel was run with the default parameters and an insert size of 461bp (6). The median insert size for the library was calculated with the CollectInsertSizeMetrics command from the Picard package. Only Pindel calls with at least 2 supporting reads, a summed mapping score greater than or equal to 100, and a mean mapping score greater than 10 were kept for further analysis. Pindel’s small insertion calls were merged if they directly overlapped, and the longest insertion length of the merged variants was used for the new variant annotation. Many of the inter-chromosomal translocations called by Pindel were located along the same regions of two chromosomes. These redundant translocation calls were merged if they spanned the same two chromosomes. All other types of Pindel calls (deletions, duplications, and inversions) were only merged if they occurred at adjacent loci.

For read pair-based structural variant calling, BreakDancer version 1.4.5 was run with the default settings (7). BreakDancer calls with a confidence score lower than 80 or with fewer than 4 supporting read pairs were removed from the analysis. The BreakDancer team reported a validation rate of 89% for indels. Additionally, insertion calls with breakpoint distances greater than the insertion size were filtered out of the final call set. Inter-chromosomal translocation calls were merged if the breakpoints from one call were both less than 1,000bp away from the breakpoints of another variant. BreakDancer’s deletions, inversions, and intra-chromosomal translocations calls were merged if they occurred at adjacent loci.

CNVnator version 0.3.2 was used to detect copy number variation along the Jurkat genome (8). The-unique flag was used for read mapping extraction, and 100bp bins were used for partitioning of the read depth signal and CNV calling. CNV calls were removed if either of the two t-test p-values were ≥ 0.01 or if the fraction of reads with mapping quality of 0 was ≥ 0.5. The raw, normalized read depth values were multiplied by 2, and integer copy number values were assigned using bins of 0.5, 1.5, 2.5, 3.5, etc.

The read depth approach is quite distinct from the other three variant calling methods due to the fact that changes in copy number can result from mechanisms that the deletion and duplication callers are unable to detect. Although the Jurkat genome is tetraploid, deletions were defined relative to a diploid genome, i.e. a copy number 1, to allow comparisons to the other methods. Duplication events, on the other hand, were defined relative to the tetraploid genome, i.e. a copy number 5.

### Variant Caller Comparisons

Structural variant calls from each tool were separated by class for further analyses. Insertions and deletions were further divided into short and long indel calls at a threshold length of 50bp.

For comparisons of insertion and deletion calls across individual software tools, two variant loci were considered to be overlapping if they had a reciprocal minimum overlap of 25%. Overlaps between duplication and inversion calls were determined less stringently. We did not require reciprocal overlap for these larger variants and called a variant site as overlapping if at least 25% of it was overlapped by another variant.

We did not calculate overlaps for homozygous deletions and inter-chromosomal translocations, as these categories of variants were compared and merged in a more involved manner.

### Variant Call Merging

Given the differences in breakpoint detection precision between the individual variant callers, the variant calls were merged hierarchically, with the more precise, split-read calls of GATK and Pindel taking precedence over the less precise variant calls from BreakDancer and CNVnator. For hierarchical merging, the GATK and/or Pindel calls were merged. Then, if present, BreakDancer calls that overlapped this merged set were removed, and the non-overlapping BreakDancer calls were merged into the GATK/Pindel set. Finally, if present, CNVnator calls that overlapped the GATK/Pindel/BreakDancer set were removed, and the non-overlapping calls were merged with the variant calls from the other tools. Loci were merged if they were either adjacent or overlapping.

Insertion calls were treated differently from the other types of structural variants. Insertion loci were only merged if they overlapped, and no hierarchy of tools was used for merging.

Homozygous deletions were also merged in a special manner. Only GATK and CNVnator are capable of making homozygous deletion calls. While GATK is able to call homozygous deletions with high precision, it can only detect relatively small deletions. On the other hand, CNVnator can identify large homozygous deletions, but its calls are much less precise. In order to minimize CNVnator’s false positive homozygous deletion calls, we removed regions with a copy number of zero if they overlapped any variants that were identified by GATK. Homozygous deletion calls from CNVnator that had greater than 50% reciprocal overlap with either Pindel or BreakDancer deletions were replaced with the Pindel or BreakDancer calls, as they identify more precise breakpoints. Finally, the filtered and swapped homozygous deletions from CNVnator were merged with the GATK calls.

### Database Comparisons

Small variant functional predictions were annotated with the SnpEff software package version 4.1 (42), and small variant database annotations were assigned with SnpSift from the same package. The latest dbSNP annotations for the hg19 reference genome were downloaded as a VCF file from the NCBI ftp repository (ftp://ftp.ncbi.nih.gov/snp/organisms/human_9606_b147_GRCh37p13/VCF/All_20160601.vcf.gz). The dbSNP file was the source of 1000Genomes information as well. The latest ClinVar annotations were also downloaded as a VCF file from NCBI (ftp://ftp.ncbi.nlm.nih.gov/pub/clinvar/vcf_GRCh37/archive/2016/clinvar_20160802.vcf.gz). ExAC data were downloaded as a VCF file from the Broad Institute (ftp://ftp.broadinstitute.org/pub/ExAC_release/release0.3.1/ExAC.r0.3.1.sites.vep.vcf.gz).

For comparisons of short insertion and deletion calls to dbSNP, 1000Genomes, and ClinVar, insertions and deletions from the VCF files that were shorter than 100bp were converted to BED format. Short insertions and deletions from dbSNP and 1000Genomes were considered a match if there was any overlap between variant loci. We required 25% reciprocal overlap between Jurkat and ClinVar variants. ClinVar records without assertion criteria were removed unless they were submitted by OMIM (Online Mendelian Inheritance in Man), as OMIM submissions are derived from literature curation. Matching ClinVar variants were then removed if they were annotated as a common variant.

The DGV database was used as a source for duplications, long insertions, and long deletions that were found in healthy individuals. DGV structural variants were downloaded as a text file from the DGV website (http://dgv.tcag.ca/dgv/docs/GRCh37_hg19_variants_2016-05-15.txt).

Long structural variants that were found by the 1000Genomes Consortium were extracted from the DGV file. Overlaps between the duplications, long insertions, and long deletions from Jurkat and DGV/1000Genomes were counted if there was greater than 25% reciprocal overlap.

Long structural variants that have been identified as pathogenic were procured from the dbVar database (downloaded from ftp://ftp.ncbi.nlm.nih.gov/pub/dbVar/data/Homo_sapiens/by_assembly/GRCh37/gvf/). All remapped and submitted germline variants for GRCh37 were downloaded, and pathogenic variants were extracted from the files. Jurkat duplications, long insertions, and long deletions were annotated as pathogenic if they had greater than 90% reciprocal overlap with a pathogenic variant from dbVar.

Affymetrix SNP6.0 array genotypes and Pindel variant calls for the Jurkat cell line (sample ID: 998184) were downloaded from the COSMIC Cell Line Project SFTP site (/files/grch37/cell_lines/v78/). Reference alleles for the SNP array data were retrieved from dbSNP, and genotype calls that either did not match with dbSNP or were homozygous for the reference allele were removed, leaving 407,817 out of 883,076 variant loci. Pindel calls from COSMIC were converted to BED format and overlaps were calculated with BEDtools, allowing for any amount of overlap between loci.

### Gene Set Enrichment

SnpEff “High Impact” variants—comprising stop gained, stop lost, start lost, splice acceptor, splice donor, exon loss, and frameshift variants—were annotated with ExAC population information. High impact variants with an ExAC minor allele frequency 0.1% were filtered out of the high impact set. The list of genes that contained a rare, high impact variant were added to the list of genes with exons that overlap a homozygous deletion, and this deleterious gene set was used to probe for biological processes that might be altered by genomic variants.

Gene set enrichment was determined using a hypergeometric test of the deleterious gene set against the MSigDB version 5.2 hallmark gene sets, canonical pathways (c2.cp), and gene ontology (GO) biological processes gene sets, using the total number of nuclear genes from hg19 (26,802 genes) as the background number of possible genes that could be drawn. MSigDB gene sets with fewer than 50 genes were not considered. MSigDB data sets were downloaded from the Broad Institute website (http://software.broadinstitute.org/gsea/downloads.jsp).

### Publication Trends

Yearly publication counts for PubMed queries were downloaded as comma-separated tables and plotted in figure 1. The following queries were used: jurkat[All Fields], jurkat[All Fields] AND apoptosis[Title/Abstract], jurkat[All Fields] AND (signaling[Title/Abstract] OR signalling[Title/Abstract]), jurkat[All Fields] AND (TCR[Title/Abstract] OR “t cell receptor”[Title/Abstract]), jurkat[All Fields] AND leukemia[Title/Abstract] OR T-ALL[Title/Abstract], and jurkat[All Fields] AND HIV[Title/Abstract].

## Acknowledgements

This work was supported by the National Institutes of Health (U19 AI063603 to D.R.S.) and by a TL1 award to L.G. through the Scripps Translational Science Institute (UL1 TR001114).

We dedicate this work to Dr. Daniel R. Salomon, who conceived of this study but passed away prior to its completion.

We thank Ali Torkamani, Luc Teyton, and Jake Bruggemann for their thoughtful comments and editing suggestions.

## References

1. U Schneider H.-U. Schwenk, G Bornkamm International journal of cancer 19, 621 (1977).

2. R. T. Abraham, A Weiss Nature Reviews Immunology 4, 301 (2004).

3. E Astoul D. A. Cantrell, C Edmunds S. G. Ward, Trends in immunology 22, 490 (2001).

4. H Li arXiv preprint arXiv:1303.3997 (2013).

5. A McKenna et al., Genome research 20, 1297 (2010).

6. K Ye M. H. Schulz, Q Long R Apweiler Z Ning Bioinformatics 25, 2865 (2009).

7. K Chen et al., Nature methods 6, 677 (2009).

8. A Abyzov A. E. Urban, M Snyder M. Ger-stein, Genome research 21, 974 (2011).

9. Jurkat, Clone E6-1 (ATCC TIB-152), https://www.atcc.org/products/all/TIB-152.aspx#characteristics. Accessed: 2017-01-30.

10. K Snow W Judd Experimental cell research 171, 389 (1987).

11. J Cheng M Haas Molecular and cellular biology 10, 5502 (1990).

12. Leibniz Institute DSMZ-German Collection of Microorganisms and Cell Lines, Cell line: JURKAT, https://www.dsmz.de/catalogues/details/culture/ACC-282.html. Accessed: 2017-01-30.

13. R Marie et al., Proceedings of the National Academy of Sciences 110, 4893 (2013).

14. K Pelak et al., PLoS genetics 6, e1001111 (2010).

15. S Levy et al., PLoS Biol 5, e254 (2007).

16. I Lappalainen et al., Nucleic acids research 41, D936 (2013).

17. S. T. Sherry, et al., Nucleic acids research 29, 308 (2001).

18. J. R. MacDonald, R Ziman R. K. Yuen, L Feuk S. W. Scherer, Nucleic acids research 42, D986 (2014).

19. S. A. Forbes, et al., Nucleic acids research 43, D805 (2015).

20. M Lek et al., Nature 536, 285 (2016).

21. A Subramanian et al., Proceedings of the National Academy of Sciences 102, 15545 (2005).

22. X Shan et al., Molecular and cellular biology 20, 6945 (2000).

23. A Sakai C Thieblemont A Wellmann E. S. Jaffe, M Raffeld Blood 92, 3410 (1998).

24. T. C. Lo, et al., Leukemia research 33, 1562 (2009).

25. T Lindsten et al., The Journal of Immunology 151, 3489 (1993).

26. M. P. Pistillo, et al., Blood 101, 202 (2003).

27. J Fargnoli et al., Journal of Biological Chemistry 270, 26533 (1995).

28. B. L. Williams, et al., Molecular and cellular biology 18, 1388 (1998).

29. S Latour L. M. Chow, A Veillette Journal of Biological Chemistry 271, 22782 (1996).

30. I Bendig N Mohr F Kramer B. H. Weber, Cancer genetics and cytogenetics 154, 22 (2004).

31. S Senturk et al., Proceedings of the National Academy of Sciences 111, E3287 (2014).

32. N. H. Shirole, et al., Elife 5, e17929 (2016).

33. R. R. Bartelt, N. Cruz-Orcutt, M Collins J. C. Houtman, PloS one 4, e5430 (2009).

34. J. P. Meijerink, et al., Blood 91, 2991 (1998).

35. M Brimmell R Mendiola J Mangion G Packham Oncogene 16 (1998).

36. L. Pérez-Carbonell, et al., Gut pp. gutjnl-2011 (2011).

37. V Piller F Piller M Fukuda Journal of Biological Chemistry 265, 9264 (1990).

38. T Ju R. D. Cummings, Proceedings of the national academy of sciences 99, 16613 (2002).

39. S Horn et al., Leukemia 18, 1839 (2004).

40. R. W. Freeburn, et al., The Journal of Immunology 169, 5441 (2002).

41. L Gioia Genomic variant data for the jurkat cell line (2017). The sequencing data used to call these variants is available at https://www.ncbi.nlm.nih.gov/sra/SRP101994.

42. P Cingolani et al., Fly 6, 80 (2012).

